# MAPK signaling links the injury response to Wnt-regulated patterning in *Hydra* regeneration

**DOI:** 10.1101/2020.07.06.189795

**Authors:** Anja Tursch, Natascha Bartsch, Thomas W. Holstein

**Affiliations:** Centre for Organismal Studies (COS), Universität Heidelberg, Heidelberg 69120, Germany

## Abstract

*Hydra* has a long history as an informative model to study pattern formation and regeneration. Wnt signaling is a critical component of *Hydra* patterning that must be activated during regeneration, but it is largely unknown how the injury stimulus ultimately leads to this activation. In a proteomic screen we previously identified mitogen protein kinases (MAPKs) among the earliest injury response factors in *Hydra* regeneration, making them attractive candidates to transmit injury-response signals to the initial steps of patterning, which in *Hydra* includes Wnt signaling. Our data demonstrate that three MAPKs, p38, c-Jun N-terminal kinases (JNKs), and extracellular signal-regulated kinases (ERK), are essential to initiate regeneration in *Hydra*. This activation occurs in response to an injury signal, which consists of calcium and reactive oxygen species (ROS) signaling. Phosphorylated MAPKs exhibit cross-talk with a mutual antagonism between the ERK1/2 pathway and the stress induced MAPKs. The activation of these MAPK pathways, as well as the induction of apoptosis, occurs in all injuries regardless of the position of the wound. MAPK phosphorylation is required for the transcriptional activation of position independent *Wnt3* and *Wnt9/10c* ligands. In summary, our data show that the activation of MAPKs is an essential component of the wound response which transmits the injury signal to induce the transcriptional activation of *Wnt* ligands, which are essential for patterning the regenerating tissue. Given the high level of evolutionary conservation of MAPKs and Wnts in the injury response, this likely represents a deeply conserved pathway in animals.

## Introduction

The ability to regenerate varies widely across species. Most vertebrates have only a limited capacity to regenerate but some animals like cnidarians and planarians can regenerate their whole body. A conserved aspect of regeneration is that signaling pathways and GRNs (gene regulatory networks) that are involved in embryonic development become re-activated during the regeneration process (Hill and Petersen, 2015, 2018; Petersen and Reddien, 2011; Tejada-Romero et al., 2015; Tewari et al., 2018; Vriz et al., 2014; Wurtzel et al., 2015). It is poorly understood, however, how the initial injury triggers pattern formation to replace lost structures in highly regenerating animals and tissues, and why this does not occur in non-regenerative animals and tissues.

We addressed the linkage between injury and patterning during regeneration in the freshwater polyp *Hydra*, a member of the ancient clade of Cnidaria, which also includes jelly fish, sea anemones and corals. *Hydra* polyps have a simple, gastrula-shaped body with a hypostome and tentacles at the oral end (the “head”) and a peduncle and pedal disc at the aboral end (the “foot”). The phenomenon of regeneration was first discovered when Trembley (1744) cut polyps into two halves. At each site of the cutting and virtually from the same gastric tissue, a head regenerated in the lower half and a foot in the upper half of the polyp (Bode, 2003; Bosch et al., 2010; Holstein et al., 2003; Trembley, 1744). On the molecular level, Wnt/7-catenin signaling is necessary and sufficient to pattern the regenerating *Hydra* head (Guder et al., 2006; Hobmayer et al., 2000; Lengfeld et al., 2009; Nakamura et al., 2011; Philipp et al., 2009; Technau et al., 2000; Vogg et al., 2019). Recent studies indicated that a transient up-regulation of 7-catenin and of several Wnts might also play an important role in *Hydra* foot regeneration (Gufler et al., 2018). Further studies have identified Erk signaling to be implicated in *Hydra* head regeneration (Arvizu et al., 2006; Chera et al., 2011; Fabila et al., 2002; Manuel et al., 2006). However, it is hitherto unknown how the Erk and Wnt signaling pathways interact during *Hydra* regeneration, and how injury-induced signaling pathways activate the patterning processes of during *Hydra* regeneration.

There is strong evidence that the injury signal contains molecular cues that are essential to induce regeneration in *Hydra*. When cutting was replaced by a careful ligature with a hair it was possible to remove the polyp’s head without injury (Guder et al., 2006; Newman, 1974). Regenerates exhibited a perfect “wound healing”, but tail pieces failed to regenerate a new head, indicating that signals released upon injury are essential to initiate this process. Studies with a head-regeneration deficient strain (*Hydra magnipapillata* reg-16) showed that after repeated wounding of the tissue the animals were able to regenerate again (Kobatake and Sugiyama, 1989; Shimizu and Sugiyama, 1993). Remarkably, a similar effect was obtained when decapitated reg-16 polyps were exposed to recombinant Wnt3 (Lengfeld et al., 2009). Taken together, these studies indicate a link between injury and Wnt expression in *Hydra* regeneration, but the molecular aspects of the injury signal that might activate Wnt signaling remained unclear.

In a comprehensive proteome and transcriptome analysis we previously identified two major phases of *Hydra* head regeneration, an early injury response and a subsequent patterning phase of the regenerating tissue that included the activation of Wnt signaling (Petersen et al., 2015). We found that several mitogen activated kinases (MAPKs) were phosphorylated upon wounding, suggesting that MAPK signaling plays an important role during the injury response (Petersen et al., 2015). Therefore, in this study we investigate the roles of these phosphorylated MAPKs, ERK1/2, p38 MAPK (p38), and c-Jun N-terminal kinase (JNK), in *Hydra* injury and regeneration. Our data show that bisecting the animal induced a cascade of wounding signals that includes Ca^2+^, ROS, MAPK signaling, apoptosis, and the activation of Wnt signaling. Importantly, an identical response occurred at both wound sites, which have different regeneration fates – one will regenerate a head and one will regenerate a foot. We also demonstrate that the upregulation of head specific genes (Wnt3, Wnt9/10c, and Wnt7) initially occurs in both head and foot regenerates. This harmonizes with a previous study that demonstrated the position independent transcriptional activation of *7-catenin* expression during the early steps of regeneration (Gufler et al., 2018). Our data show that MAPKs are an essential part of the wound response that is required for the injury signal to activate patterning. We propose a model in which differential Wnt activation is encoded by ATF transcription factors that are regulated by MAPK signaling. The ensuing fine-tuned cascade of Wnt expression is subsequently required for pre-patterning of the lost body part. Given the high level of evolutionary conservation of MAPKs and Wnts, this signaling network may be a fundamental feature of the animal wound response.

## Results

### MAPK activation is a general response to injury in Hydra

Previous phospho-proteome analysis in *Hydra* revealed rapid changes in protein phosphorylation and dephosphorylation in response to injury (Petersen et al., 2015). Among other factors, several MAPKs showed a rapid enrichment of phosphorylation upon wounding and thus may play an important role during the injury response. Because of their sharp increase in phosphorylation, we selected ERK1/2, p38, and JNK for biochemical analysis to evaluate their phosphorylation dynamics upon injury.

First, we analyzed how the levels of ERK1/2, p38, and JNK phosphorylation change in response to injury and regeneration. We used antibodies specific for activated (phosphorylated) MAPKs and analyzed animals wounded by three horizontal cuts into the body column. At various time points after injury, we prepared whole tissue lysates for Western blot analysis (Fig.1B,S1). All three MAPK proteins showed an increase in phosphorylation within the first minute post injury (pi), and maximum phosphorylation levels were reached at 30 minutes pi. Interestingly, while the phosphorylated JNK and p38 levels started to decrease after 60 minutes pi, phosphorylated ERK was still detectable until 6 hours pi (Fig. S1A). The observed increase in MAPK phosphorylation levels was not due to an increase in total protein concentration (Fig. S1C). We found a similar activation of MAPKs in regenerating head and foot pieces when we bisected the animals with a scalpel to create wounds that will regenerate (Figure 1B).

**Figure 1.**
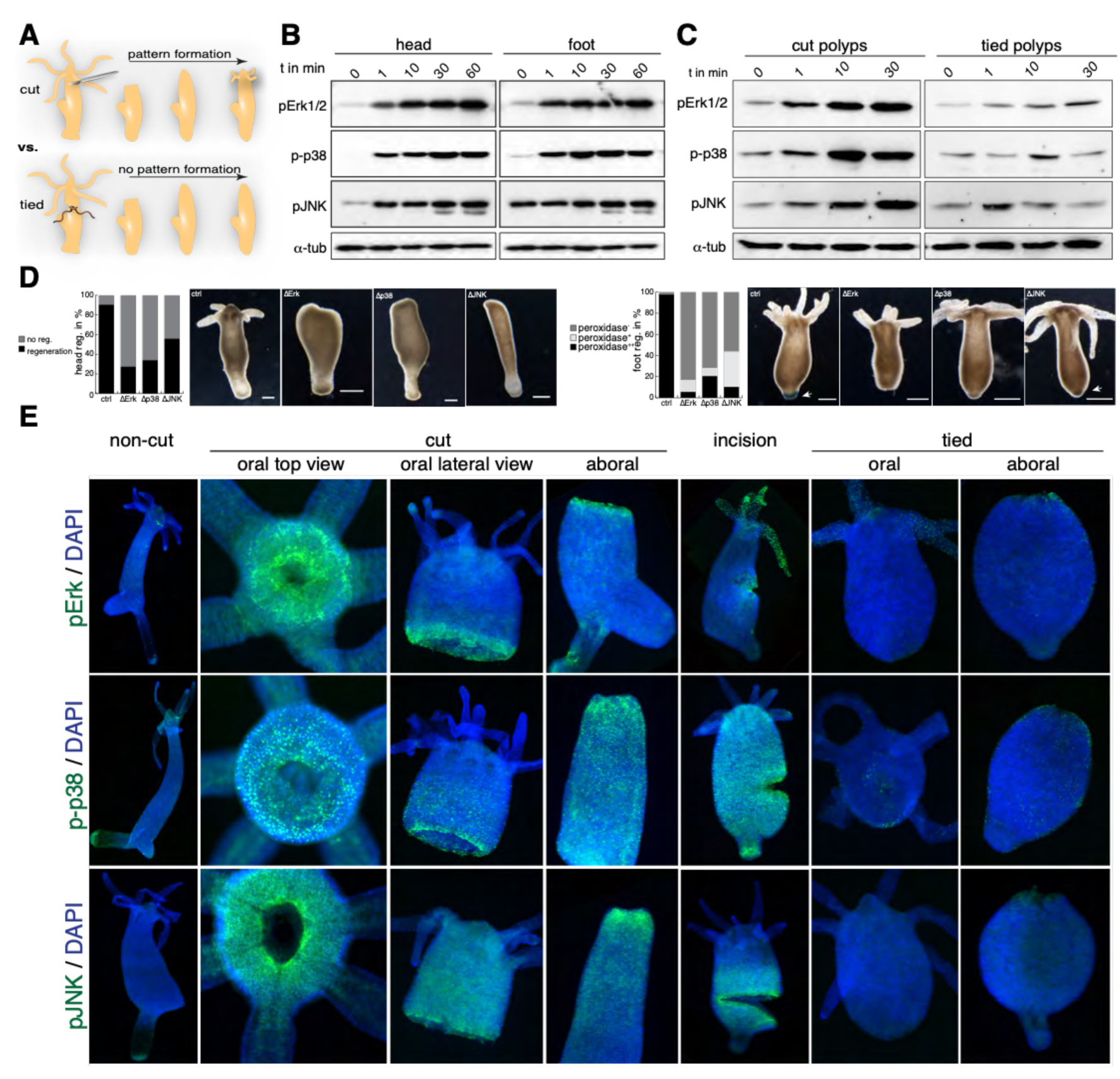
MAPKs are activated in an injury dependent manner. **A)** Scheme representing successful pattern formation upon injury, whereas removal of the head without wound response fails a proper regeneration. **B)** Proteome and phospho-proteome analysis reveal ERK1/2, p38, and JNK as potential candidates to be enriched upon injury. **C)** Western blot analysis of phosphorylated ERK 1/2, p38, and JNK. Head and foot regenerates showed MAPK activation to similar extent. **D)** Western blot analysis of phosphorylated (activated) levels of ERK1/2, p38, and JNK. Whereas wounded animals showed an increase over time, tied animals exhibited only poor signals of activated MAPKs. **E)** Regeneration assay of polyps inhibited for either ERK1/2, p38 or JNK. Inhibition of ERK1/2 via MEK1/2, p38 and JNK reduce the regeneration capacity of both, foot and head. Foot regeneration was visualized by performing a peroxidase stain. Head regeneration: control n=108; ΔErk n=61; Δp38 n=58; ΔJNK n=72, foot regeneration n=45 per condition. **F)** Non-cut animals only show a weak signal of activated MAPKs (green). In contrast, cut animals show a strong signal at the wounded edge, which is distributed in a gradient-like manner along the body column. Activated levels of MAPKs were detectable irrespective of position and severity (partial cut) of the wound. In contrast, tied animals show poor signals of phosphorylated ERK1/2, p38, and JNK.

Next, we used immunofluorescence analysis to determine the localization of pERK, pJNK and pp38 relative to the wound site. Activated MAPKs were distributed in a gradient emanating from the wounded edge (Fig.1E, cut) in both wounds that will regenerate a head and wounds that will regenerate a foot. Whereas ERK and JNK localization appears cytoplasmic, p38 expression appears nuclear (Fig.S1B). We also found a similar pattern of MAPK activation when making a partial cut on the side of the body, which is a non-regenerative injury (Fig.1E, partial cut). Taken together, our data support the conclusion that phosphorylation of JNK, p38, and ERK is a general response to injury that occurs even when regeneration is not induced.

To test whether the activation of the MAPK directly depends on the injury signal, we also bisected the animals by carefully tying off the head without wounding (Guder et al., 2006; Newman, 1974). Compared to animals cut with a scalpel, the levels of phosphorylated ERK1/2, p38, and JNK were much lower (Fig. 1C). This was also consistent with our immunofluorescence data. Compared to animals cut with a scalpel, the levels of phosphorylated ERK1/2, p38, and JNK were much lower (Fig. 1E), which demonstrated that MAPK phosphorylation at the onset of regeneration was dependent on the injury signal. These findings implicate MAPK activation as a critical part of the injury signal that may be important for regeneration to occur.

### MAPK activation is required for regeneration

Given their activation in response to injury, we next tested if MAPKs are required for head and foot regeneration using inhibitors specific for ERK1/2 (U1026), p38 (SB203580), and JNK (SP600125). U0126 is an inhibitor of the mitogen-activated protein kinase kinase family members MEK 1/2 in the MAPK pathway, which are the upstream kinases of ERK1/2. However, due to its specificity, we will refer to it as an ERK1/2 inhibitor for simplicity. To test the effect of these inhibitors on head regeneration, heads were removed by cutting at a 70% distance from the aboral end and the fraction of properly regenerated heads at 72 hours post-amputation was determined (Fig. 1D). The majority of control animals (92%) exhibited a fully regenerated head with a hypostome and a ring of tentacles. By contrast, animals treated with MAPK inhibitors showed regeneration deficiencies. Although the wound was closed, no sign of head regeneration was found in 72%, 66% and 44% of the animals treated with ERK, p38 and JNK inhibitors, respectively. To test the function of MAPKs during foot regeneration, we dissected animals at 30% of the body length and used the peroxidase assay to visualize differentiated foot tissue (Hoffmeister and Schaller, 1985). In untreated controls, 98% of *Hydra* were positive for peroxidase activity at the aboral end 72 hours post amputation (hpa), which indicates full regeneration of the basal disc. By contrast the majority of inhibitor-treated animals did not show a similar level of peroxidase activity staining (Fig. 1D). Our data thus clearly show that the early activation of MAPKs by an injury signal is essential for regeneration to proceed normally, regardless of the type of the tissue to be regenerated.

### MAPK activation depends on ROS signaling

Given the importance of MAPK signaling in regeneration, we next set out to determine how MAPKs are activated in response to injury. We considered reactive oxygen species (ROS) because recent work in zebrafish and *Drosophila* showed the importance of ROS for regeneration, probably by either activating or modulating different signaling pathways such as JNK or Hedgehog signaling (Gauron et al., 2013; Romero et al., 2018). To test whether ROS is produced at the site of injury, animals were incubated with CellRox, a ROS sensitive dye, prior to amputation (Fig. S2). Indeed, bisected animals incubated with this dye showed a fluorescent signal at the wounded edge indicating ROS production (Fig. S2A). The signal was abolished when animals were additionally incubated with reduced glutathione (GSH) or bisected without injury by tying (Fig. S2A, lower panel). To test whether the redox state has an impact on the activation levels of MAPKs, animals were incubated in *Hydra* medium (HM), or in HM supplemented with either hydrogen peroxide (H_2_O_2_) or GSH to produce an oxidizing or reducing environment, respectively. The exposure to H_2_O_2_ resulted in elevated activation levels for all MAPKs tested (Fig. 2A). The reducing environment did not result in any detectable activation of p38 and JNK upon wounding and the phosphorylation of ERK was diminished (Fig. 2A). These findings were further supported by immunofluorescence analyses using the same experimental setup. In the presence of H_2_O_2,_ the signal intensity of activated ERK, p38, and JNK was increased and their distribution along the tissue was expanded (Fig. 2B, S2C). Treatment with GSH reduced levels of p38 and JNK activation throughout the tissue, while phosphorylated ERK became restricted only to the wound edge (Fig. 2B).

**Figure 2.**
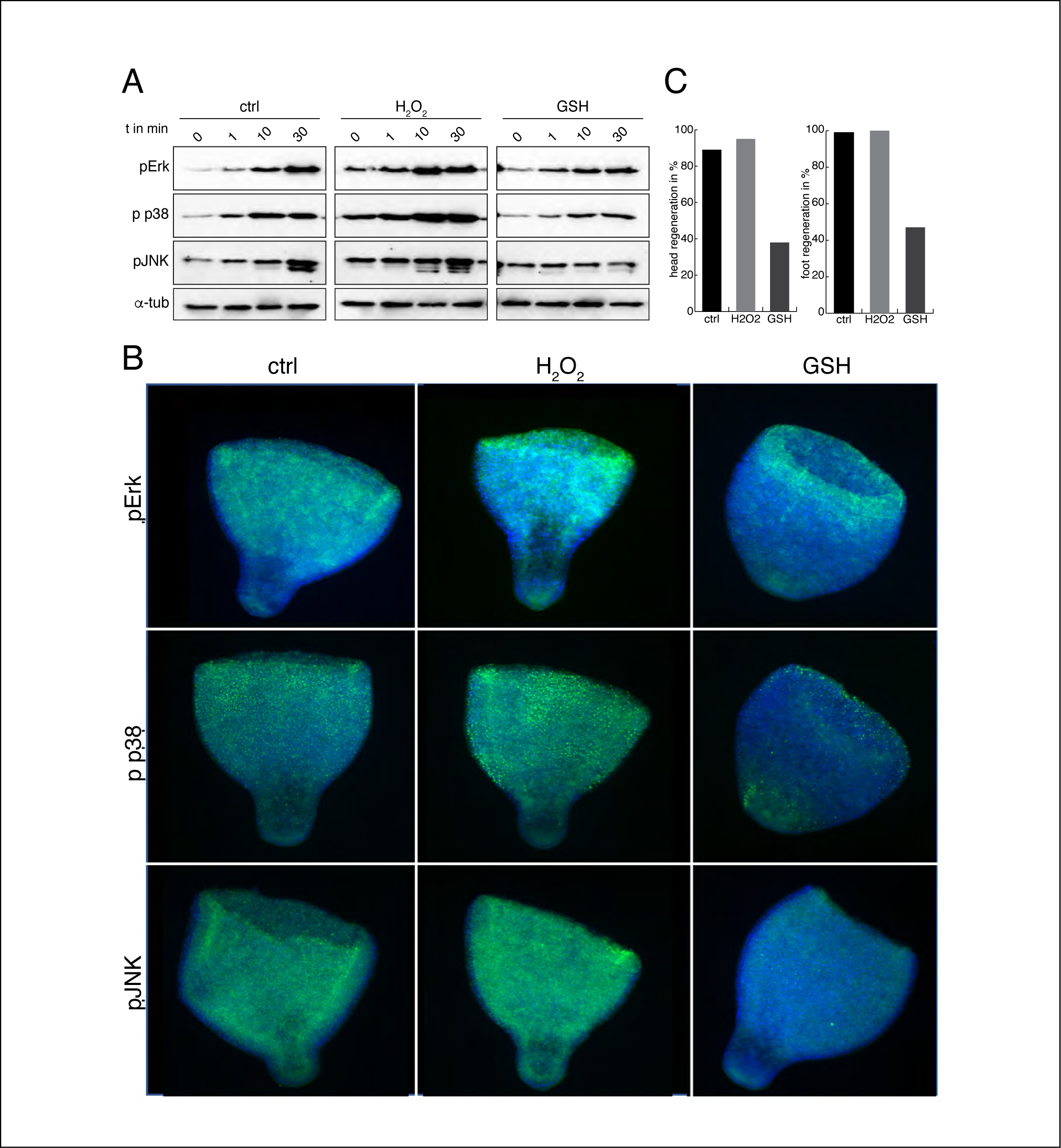
MAPKs are activated by ROS signaling. **A)** Western blot analysis of activated ERK1/2, p38, and JNK. Whereas exposure of *Hydra* to H_2_O_2_ results in elevated activation levels, exposure to reduced GSH resulted in reduced levels of phosphorylated MAPKs. **B)** Immunofluorescence analysis of activated MAPKs upon treatment with H_2_O_2_ or GSH. Exposure to H_2_O_2_ resulted in an increase of the signal intensity all over the body column. In contrast treatment with GSH resulted in decreased activated levels particularly for p38 and JNK. **C)** Evaluation of head and foot regeneration 72 dpa (n=40 each). Regeneration capacity did not significantly improve in an oxidizing environment (H_2_O_2_), but dropped in the presence of GSH.

We next tested if manipulating the redox state has an effect on regeneration by incubating bisected *Hydra* in the presence of either H_2_O_2_ or GSH (Fig. 2C; S2B). Animals regenerated normally under oxidative conditions. By contrast, 64% of *Hydra* head regenerates and 47% of *Hydra* foot regenerates did not proceed under reducing conditions (Figs. 2C, S2B). These data support a model in which ROS production in response to injury triggers the activation of MAPKs which is required to promote regeneration in *Hydra*.

### MAPK activation depends on Ca2+ signaling

Calcium signaling is one of the fastest transduction pathways and it is involved in many processes including wound closure and regeneration (Clapham, 2007; Xu and Chisholm, 2011; Yoo et al., 2012). Given the fast kinetics of MAPK phosphorylation in response to wounding (Fig. 1; S1), we hypothesized that Ca^2+^ signaling is responsible for the gradient-like transmission of the injury signal and activation of MAPKs. To test this, we increased the intracellular calcium levels by treating polyps with the Ca^2+^ ionophore A23187. We observed that increased cytoplasmic Ca^2+^ levels during regeneration resulted in elevated phosphorylation levels of ERK, p38, and JNK (Fig. 3A), while incubation with BAPTA, a chelating agent specific to Ca^2+^, caused a reduction in the levels of activated MAPKs compared to untreated control animals (Figs. 3A). Consistent with this, when we analyzed the effects of different Ca^2+^ levels in wounded animals *in situ*, it was evident that after BAPTA treatment, activated MAPKs became strongly reduced at the site of cutting. By contrast, A23187 treatment lead to an upregulation of activated MAPKs along the entire body column (Fig. 3B).

**Figure 3.**
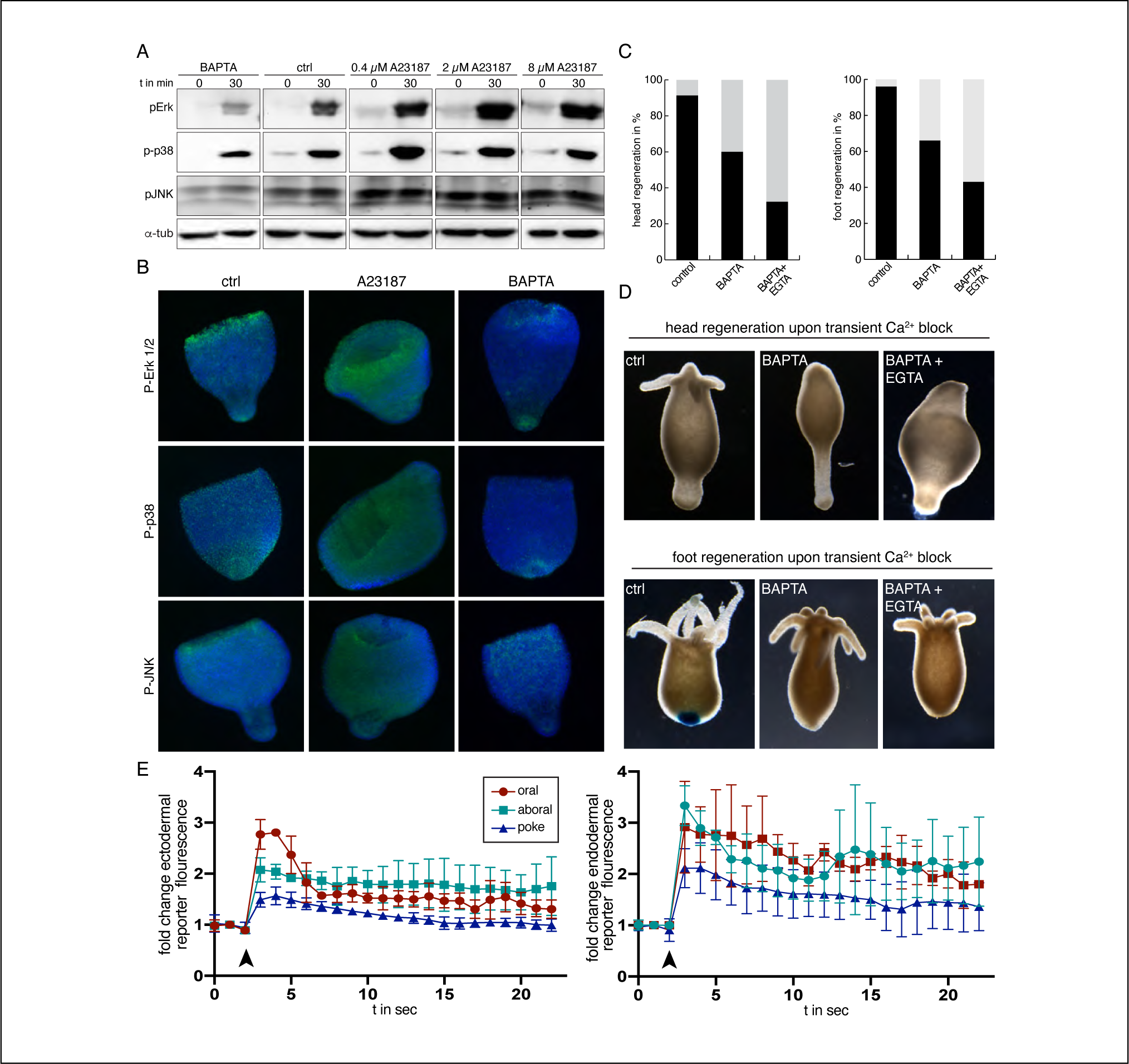
MAPKs are activated in a calcium dependent manner. **A)** Western blot analysis of activated MAPKs upon calcium concentration manipulation. Reduction of intracellular calcium levels using BAPTA resulted in a drop of activated MAPK levels. Titration of the ionophore A23187 induced increased phosphorylation events of ERK1/2, p38, and JNK. **B)** Representative immunofluorescence pictures of polyps treated with either A23187 or BAPTA. Increase of cytoplasmic calcium levels turned gradient-like distribution of activated MAPKS (ctrl) into a homogenous stain all over the tissue (A23187). In contrast, calcium capture by BAPTA yielded in a reduction of signal intensity upon injury. **C)** Quantitative analysis of regeneration capacity upon transient calcium reduction for up to 3 hours post injury. A transient repression of calcium signaling by BAPTA reduced head and foot regeneration (n=40 each). Regeneration capacity was further decreased when BAPTA was used in combination with EGTA. **D)** Representative pictures of quantitative data shown in C. **E)** Fold change of GCaMP reporter fluorescence from ectodermal (left) and endodermal (right) transgenic strains upon simple contraction (poke) and injury (cut). Moment of poking or cutting the polyp is indicated by black arrowhead. The slightly higher reporter signal intensity in cut animals upon injury as well as the increased duration of calcium release was detectable irrespective of the positional value (oral, aboral) and germ layer (ectoderm, endoderm).

We next tested the effect of Ca^2+^ levels on regeneration capacity. We found that reducing Ca^2+^ for the first 3 hours after wounding with BAPTA treatment reduced the level of successful head regeneration to 60% and a co-treatment with EGTA resulted in a further reduction to 32% (Fig. 3C). Likewise, the level of successful foot regeneration dropped to 65% in the presence of BAPTA and to 45% when a combination of BAPTA and EGTA was used.

Since calcium is not exclusively released upon injury, but is also released during muscular contractions, we next asked if the injury signal produces a Ca^2+^ signal pattern that is distinct from the muscle contraction. By using transgenic animals expressing a genetically encoded Ca^2+^ indicator (GCaMP) in the ectodermal or endodermal epithelial cell lineage, we were able to measure cellular Ca^2+^ levels in polyps that were bisected by a cut (injury) or by poking (no injury). We found that both the amplitude and the duration of the Ca^2+^ pulse were increased upon injury (Fig. 3E) (movie S1-4). This effect was also observed with calcium-free *Hydra* medium, which excludes the possibility of reporter activation due to an artificial Ca^2+^ influx from the medium (Fig. S3, movie S5-8). The amplitude of the Ca^2+^ release was lower in the endodermal cells, but otherwise, no difference was found in the dynamics (frequency, amplitude width) of the Ca^2+^ waves between the two epithelial layers. We therefore assume that both epithelia contribute equally to injury-induced calcium signaling. Thus, injury promotes both the release of ROS and Ca^2+^ that in turn acts positively on the activation of MAPKs.

### Apoptosis exhibits position independent activation during regeneration

Previous studies have indicated that apoptosis is indispensable for inducing wound healing and regeneration (Fogarty and Bergmann, 2017) (Chera et al., 2009). To test whether there is a link between MAPK activation and induction of apoptosis, we studied apoptosis using the classical TUNEL staining protocol (see Materials and Methods). TUNEL assays were performed on both control or MAPK inhibited animals. To our surprise (Chera et al., 2009), the apoptotic cells were not restricted to head regenerating animals, but were detectable to the same extent in tissue of foot regenerating animals (Fig. 4). To assess the temporal progression of apoptotic cells we analyzed four different time points during early head and foot regeneration (0.5, 1, 2, and 4 hpa). Apoptosis can be detected as early as 30 minutes after amputation and reaches a maximum at 1-2 hpa (Fig. S4A). TUNEL positive cells coincided with DAPI positive nuclei and apoptotic cells were located close to the amputation site, whereas the remaining tissue was almost free of any signal. The number of TUNEL positive apoptotic cells dropped again by 4 hpa. We also tested the position dependency of apoptosis and bisected polyps at 80% and 20% body length. Strikingly, animals cut at 80% and 20% body length showed the same kinetics of apoptotic cells (Fig. S4B). This indicates that apoptosis occurs independently of the cut height and is a general event in both head and foot regenerates.

**Figure 4.**
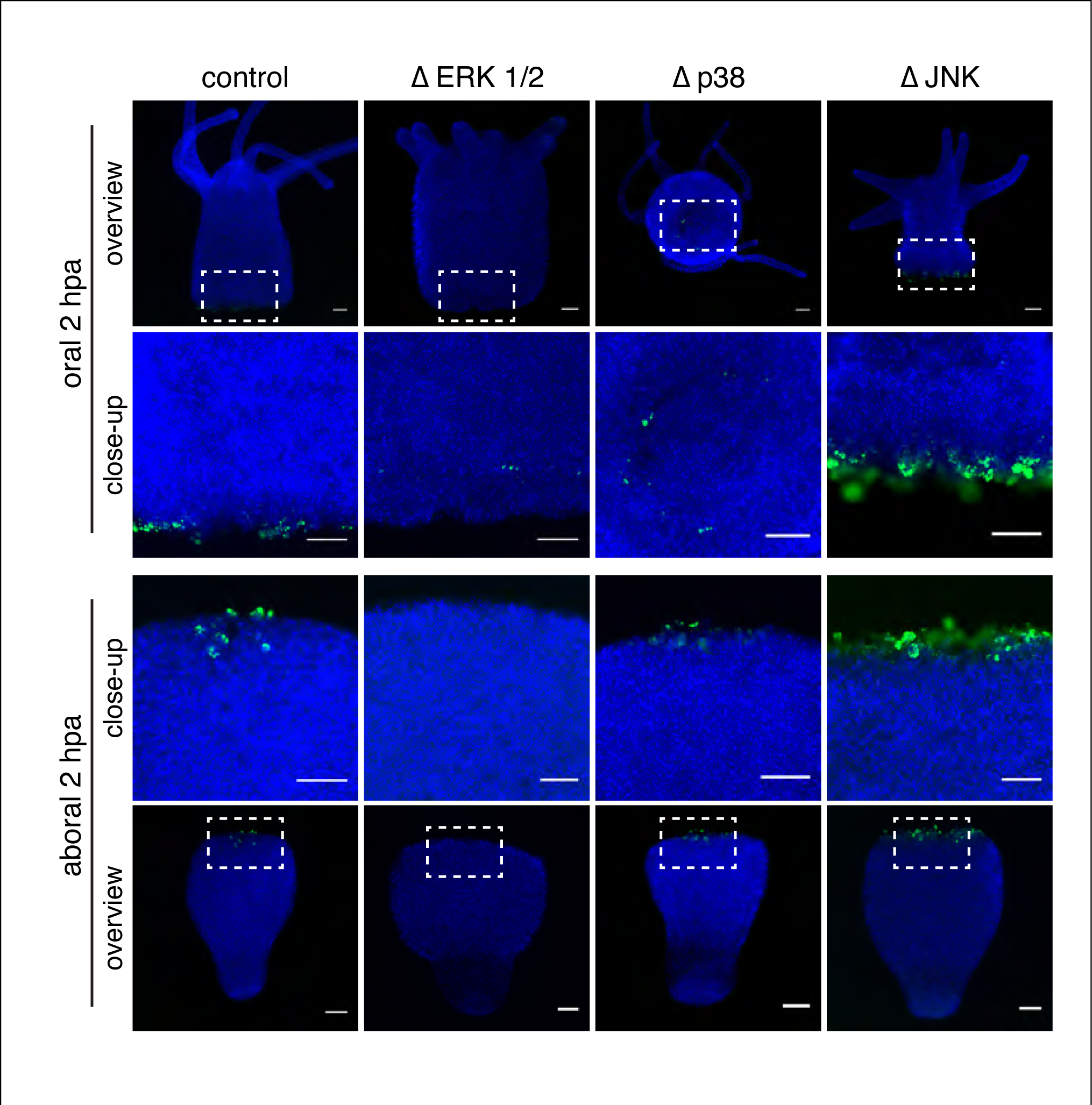
MAPKs influence frequency of apoptosis upon injury. TUNEL assays performed at 2 hpa possess apoptotic events at both the head and the foot regenerate (ctrl). Interestingly, pharmacological inhibition of Erk (ΔErk) yields in absence of apoptotic cells, while inhibition of JNK (ΔJNK) caused a strong increase of apoptotic events, while diminished p38 activity showed no effect compared to the control. Scale represents 100µm.

Since the importance of the interstitial cell lineage during regeneration has been much debated in many studies over the last years we also analyzed the cell type specificity of apoptosis by using animals that were depleted of interstitial stem cells after hydroxyurea (HU) treatment or by using heat-shocked animals of the SF-1 mutant strain (Fujisawa, 2003). After bisection at 50% of such epithelial animals, the kinetics, localization and number of apoptotic cells that appeared in the regenerates was the same as in untreated controls with regard to sf-1 polyps, while treatment with HU resulted in even higher numbers of TUNEL positive cells (Fig. S4C). This result demonstrates that epithelial cells undergo apoptosis in both head and foot regenerates.

Remarkably, the number of TUNEL-positive cells increased upon inhibition of the JNK pathway, which is in line with the previous observation that a transient activation of JNK in primary murine embryonic fibroblasts by the tumor necrosis factor (TNF) results in cell survival (Ventura et al., 2006). In contrast, a decrease of apoptotic cells was detectable upon ERK1/2 inhibition (Fig. 4) while reduced p38 activity did not show any significant difference compared to control conditions (Fig. 4). Overall, these results indicate that apoptosis is part of a general injury response rather than being an event specific to head regenerates.

### ERK1/2, p38, and JNK pathways participate in extensive crosstalk

MAPKs are frequently activated in parallel (Cargnello and Roux, 2011) suggesting that the ERK, p38, and JNK pathways are integrated by positive and negative feedback, which modulates the signal in specific ways to trigger different cellular fates (Fey et al., 2012). The effects of MAPKs on apoptosis suggest that ERK1/2 acts antagonistically to the stress-inducible MAPKs p38 and JNK. To explore these dynamics during regeneration, we tested feedback interactions between the ERK1/2, p38, and JNK pathways using pathway-specific inhibitors. Animals were incubated for one hour with the inhibitors of these pathways, bisected, and then processed at different time points for Western blot and immunofluorescence analysis. As expected, Western blot analysis revealed an almost complete inhibition of phosphorylated ERK in the presence of the ERK inhibitor (U0126) (Figs. 5A, S5). Phosphorylated ERK was also inhibited when we pretreated the animals with the JNK inhibitor (SP600125), but was activated in response to treatment with the p38 inhibitor (SB203580). Conversely, activated levels of p38 increased upon ERK inhibition, but decreased in the presence of the JNK inhibitor. By contrast, phosphorylation of JNK was reduced after treating the polyps with p38 inhibitor, but elevated upon inhibition of ERK (Fig. 5A, B).

**Figure 5.**
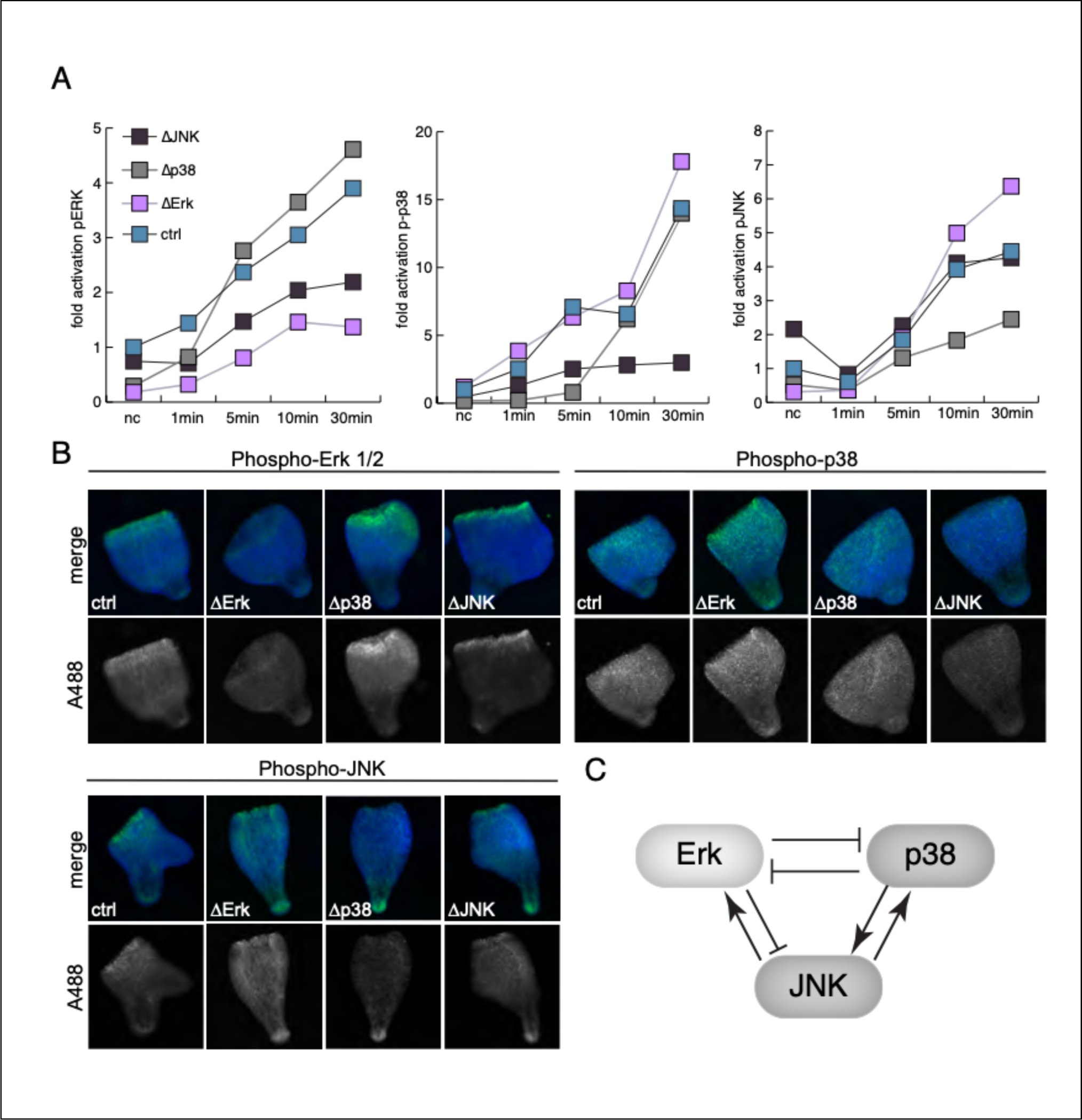
MAPKs underlie extensive crosstalk. **A)** Quantitation of activated MAPKs Western blot analysis upon inhibition of either ERK1/2 via MEK1/2, p38, and JNK. **B)** Representative pictures of phosphorylated MAPK upon inhibitor treatments. **C)** Scheme showing the relationships among MAPK pathways based on data from A and B. Whereas stress-activated MAPKs (p38 and JNK) activate the respective pathway, the ERK pathway inhibits stress-activated MAPKs.

These findings were also reflected by immunofluorescence analysis (Fig. 5B). It is important to note that the interference with a particular pathway only changed the relative phosphorylation levels of the respective pathways, while the distribution pattern itself remained unchanged (Fig 5B). In summary, these data indicate a positive feedback between the stress-inducible p38 and JNK pathways. In contrast, the ERK1/2 pathway inhibits the stress-inducible pathways, which is in line with previous findings (Fey et al., 2012). Strikingly, there is a mutual antagonism between the p38 and ERK1/2 pathways, those that showed severest regeneration phenotypes upon inhibition.

### A MAPK network modulates Wnt signaling

Wnt expression is essential for successful regeneration in *Hydra* (Hobmayer et al., 2000). Upon decapitation, several Wnt ligands are transcriptionally activated in a temporal sequence and participate in patterning the regenerating *Hydra* head (Lengfeld et al., 2009). Strikingly, recent results have also shown that a transient up-regulation of β-catenin occurs early during both head and foot regeneration and treatment with a Wnt pathway inhibitor blocks regeneration of both structures (Gufler et al., 2018). Whereas β-catenin expression is maintained in the presumptive head, it is subsequently downregulated in the presumptive foot tissue. These data suggest that β-catenin, the downstream target of canonical Wnt signaling, is initially part of a generic response to wounding.

Given our results that inhibition of MAPK activation blocked both head and foot regeneration, we next asked how the activation of either ERK, p38 or JNK affects early *Wnt* expression. Thus, animals were bisected at 50 % body length and the expression levels of *Wnt3, Wnt9/10c*, and *Wnt7* were evaluated by qRT-PCR in controls and upon MAPK inhibition at different time points post amputation. Previous in situ hybridization studies revealed that *Hydra Wnt9/10c* and *Wnt3* were detected within the first three hours of head regeneration while *Wnt7* expression was observed starting at 12 hours (Lengfeld et al., 2009). In this study, using qRT-PCR we observed a strong upregulation of *Wnt 9/10c* and *Wnt3* at 3 hrs in both head and foot regenerates, while the upregulation of *Wnt7* at this time point was much smaller (Fig. 6A). Upon ERK1/2 inhibition, all *Wnt* genes were down-regulated with the exception of *Wnt3* in foot regenerates, suggesting the ERK1/2 pathway activates Wnt expression either directly or indirectly. Upon inhibition of the p38 or JNK pathways, which are the stress-induced MAPK pathways, we found an upregulation of *Wnt3* and *Wnt7* by a factor of 6-20 in both head and foot regenerates (Fig. 6B, C). By comparison, in head and foot regenerates the expression *Wnt9/10c* was downregulated (Fig. 6B, C). Thus, p38 and JNK have a contrasting effect on *Wnt9/10c* and *Wnt3*. Since animals that were inhibited for p38 or JNK do not show any signs of regeneration despite the increased expression levels of *Wnt3* and *Wnt7*, we propose that *Wnt9/10c* acts as the primary *Wnt* gene in the *Wnt* gene cascade during regeneration.

**Figure 6.**
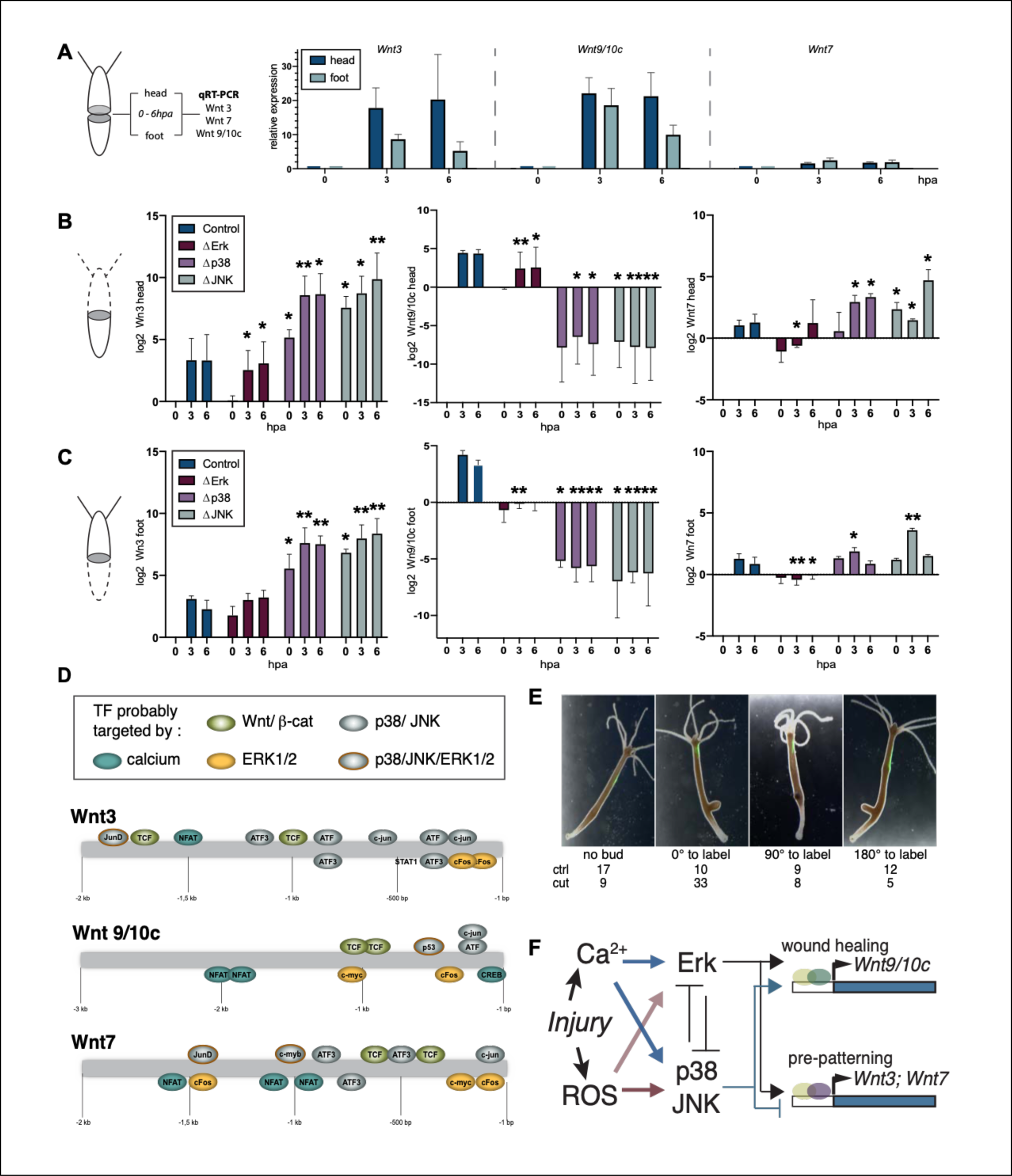
MAPKs influence Wnt expression levels. **A)** Fold change expression levels of Wnt3, Wnt9/10c, and Wnt7 in head and foot regenerates upon injury. **B-C)** Log2fold change of Wnt genes in head and foot regenerates upon pharmacological MAPK inhibition. While abrogated Erk activity generally resulted in reduced *Wnt* expression levels, inhibition of stress induced MAPKs yielded in higher expression levels for *Wnt3* and *Wnt7*, but in a down-regulation for *Wnt9/10c*. *p-value<0.05; **p-value<0.005. Bars without asterisk did not pass significance threshold. **D)** Schematic representation of promotor analysis in order to identify transcription factors potentially regulated by MAPKs. While motifs of ATF3 were obtained for both, Wnt3 and Wnt7, only one ATF motif was predicted for the Wnt9/10c promotor. **E)** Polyps were labeled with GFP latex beads and 2 h post labeling either incised in the budding region at the side of the label (cut) or left remain unaffected (non-cut). Evaluation of bud development was performed 3 days post -labeling. Negative regression analysis revealed a probability being 5.7 times higher to develop a bud at the site of label, when *Hydra* was incised (p>0.0001). **F)** Proposed model of a position-independent wound response. Injury causes a release of ROS and calcium that in turn affect phosphorylation state of MAPKs. The limited effect of ROS on Erk is indicated by less opaque arrow. Erk and stress inducible MAPKs exhibit an antagonistic relationship that tis also reflected on the transcriptional level of *Wnt3/Wnt7* and *Wnt9/10c*.

We next asked how this differential *Wnt* expression profile upon MAPK might be achieved at the transcriptional level. Among the important regulators of wound healing in metazoans are activator protein-1 (AP-1) leucine zipper transcription factors, that are composed of hetero-or homodimers of the Fos (e.g. c-Fos), Jun (e.g. c-Jun), and CREB/ATF protein family members (Schafer and Werner, 2007). We performed an *in-silico* prediction of transcription factor binding sites for the *Wnt* promotors and subsequently screened for transcription factors likely regulated by MAPKs. For *Wnt 9/10c* we found a consensus sequence with high probability scores that is likely to be bound by the Activating Transcription Factors (ATFs) (Fig. 6D). This metazoan-specific protein family has been frequently reported to be phosphorylated and thus activated by the JNK and p38 pathway (Lopez-Bergami et al., 2010; Ouwens et al., 2002; Wang et al., 2019; Watson et al., 2017). Whereas the *Wnt9/10c* promotor had higher probability scores for ATF and ATF2/CREB, which act as activating transcription factors, the promotors of *Wnt3* and *Wnt7* had a higher binding probability for ATF3, which is known to act as a repressor (Fig. 6D) (Hai, 2006; Hai et al., 1999; Inoue et al., 2018; Katoh, 2018; Zhou et al., 2017). Interestingly, one consensus sequence of ATF3 was further predicted to be located in the repressor element of *Hydra Wnt3* (Nakamura et al., 2011). Since ATF3 is closely related to Fos, while ATF2 shows higher similarity to the ATF7 and Jun family, alignments and subsequent phylogenetic analysis has been executed to validate the existence of the *Hydra* homologues (Fig. S6A-B) (Vinson et al., 2002). We presume that ATF transcription factors bind the promotor region of Wnt 9/10c and Wnt 3 and are responsible for the antagonistic effects of MAPKs on different Wnts (see also Cazet & Juliano, 2020).

Since asexual reproduction in *Hydra* likewise depends on Wnt expression, we also asked whether we are capable of inducing budding by incising in the budding region of *Hydra* to test our hypothesis that the injury response activates *Wnt* expression. Thus, polyps were labelled with fluorescent microspheres (Fig.6F). Two hours post-labeling one population remained unaffected, while the other group was incised in the budding region at the labelled site. Remarkably, while 20% of non-cut polyps developed a bud at the side of labeling, the frequency increased to 60% when animals have been incised (Fig.6F). Logistic regression analysis revealed this outcome to be significant (p-value<0.0001) with a probability of biased budding being 5.7 times higher upon induction of an injury compared to non-incised polyps (Fig.6F). This result demonstrates that an injury directly activates *Wnt* expression. However, it also indicates the requirement of a competent tissue capable to respond to the stimulus, since incision at any other position beyond the budding zone did not show an effect.

## Discussion

*Hydra* has one of the most outstanding regeneration capacities in the animal kingdom, ranging from the simple reconstruction of lost parts to *de novo* pattern formation from cell aggregates. This ability is critically dependent on a cascade-like activation of Wnts that initiate regeneration and orchestrate particular stages of patterning (Hobmayer et al., 2000; Lengfeld et al., 2009). Based on a proteome and transcriptome analysis (Petersen et al., 2015), which showed that MAPKs are quickly activated after an injury, we can show now that a MAPK network is required for the activation of the Wnt expression cascade during regeneration (see Fig. 6F for summary). We demonstrate that the injury produces Ca^2+^ and ROS signals that in turn activates ERK1/2, JNK, and p38 in a <u>position-independent</u> manner. Our work identified stress-induced MAPKs that antagonize ERK function and balance apoptosis and Wnt signaling. Remarkably, our data suggest that the antagonism on the expression level is mediated by ATFs which includes ATF3, a hitherto little-known transcription factor in *Hydra* that is activated upon stress stimuli and induces the complex GRN of *Wnt* gene activation.

### The injury stimulus

Aberrant or missing injury signals can lead to a complete loss of regenerative capacity (Guder et al., 2006; Newman, 1974; Yoo et al., 2012). Our data on *Hydra* show that the injury signal is generated quickly and leads to the phosphorylation of MAPKs. This fast generation seems to be crucial to initiate wound healing. We were able to demonstrate that the injury signal is composed of Ca^2+^ and the production of ROS, which has not been studied in early *Hydra* regenerates so far (Galliot et al., 2017)(Gufler et al., 2018). Both signals transmit the information about tissue damage intracellularly by activating ERK1/2, JNK, and p38. The activation by ROS is likely achieved by inactivation of counteracting protein tyrosine phosphatases via oxidation of regulatory cysteines, while calcium has been frequently reported to activate MAPK via calmodulin dependent kinases, PKC and Ras activation (Chuderland and Seger, 2008; Farnsworth et al., 1995; Katz et al., 2006; Paulsen and Carroll, 2010). Although there is increasing evidence for the importance of either ROS or calcium signaling upon injury, it is unclear whether ROS production is Ca^2+^ mediated or whether both messengers act independently in the regenerative context as it was shown for zebrafish fin regeneration (Yoo et al., 2012).

The phosphorylation of MAPKs was also among the earliest regeneration responses in our proteomic/transcriptomic study on *Hydra* head regeneration (Petersen et al., 2015). Here we show that the activation of ERK1/2, p38 and JNK is essential for both head regeneration and foot regeneration, with ERK1/2 showing the most sustained activation kinetics (Fig. S1). Since MAPK phosphorylation was observed upon both incision and amputation irrespective of their localization, it implicates MAPKs as part of a generic wound response. The term “generic wound response” was initially introduced by Reddien and his colleagues, who were able to show that both incision and amputation promote a position-independent up-regulation of the same set of genes in planarians (Wenemoser et al., 2012; Wurtzel et al., 2015). This finding was further specified by Owlarn & Bartscherer who showed a sustained increase of phosphorylated ERK in response to incision and amputation, which was required for Wnt activation (Owlarn et al., 2017). In contrast, the role of stress-induced MAPK pathways in regeneration is only poorly investigated. A joint activation of both stress inducible MAPK pathways was so far only reported for axonal regeneration in *C. elegans* and imaginal disc regeneration in *Drosophila* (Nix et al., 2011; Santabarbara-Ruiz et al., 2015), while most studies either focus on ERK activation during regeneration (Makanae et al., 2013; Owlarn et al., 2017; Suzuki et al., 2007; Yun et al., 2014) or p38/JNK (Chen et al., 2007; Tejada-Romero et al., 2015), our data indicate that these pathways do not work independently of one another, but participate in an extensive cross-talk. Inhibitor experiments revealed that p38/JNK interacts with Erk1/2 as potential antagonist. It will therefore be interesting to see whether this antagonism also holds true for other regeneration-competent organisms for which the critical role of Erk activation has been already demonstrated during regeneration.

A major part of the wound response is the caspase-3 mediated activation of apoptosis, which has been linked to the stimulation of proliferation in planarians, newts, and flies and it was referred to as apoptosis-induced proliferation (AiP) (Fogarty and Bergmann, 2017; Hwang et al., 2004; Ryoo et al., 2004; Vlaskalin et al., 2004). The mechanisms are unclear so far, but several context- and tissue-specific forms of AiP have been described where mitogens are secreted in parallel or independently to the final execution of apoptotic cells (Fogarty and Bergmann, 2017). Initiators of caspase-dependent AiP could be ROS and JNK signaling, whereas the latter has been already described to directly promote *Wnt* and *TGF-beta* expression via a hitherto unknown mechanism in *Drosophila* (Diwanji and Bergmann, 2018; Fan et al., 2014; Fogarty and Bergmann, 2017; Fogarty et al., 2016; Perez-Garijo et al., 2009).

In *Hydra*, Galliot and coworkers have proposed a model in which caspase-3 activation of apoptotic interstitial cells promotes the secretion of Wnt3 which in turn activates the proliferation of epithelial cells (Chera et al., 2009; Fogarty and Bergmann, 2017). This activation was also described to be head-specific (Chera et al., 2009). While this is a highly elegant hypothesis for generating positional information, it could not be confirmed by our experiments, since none of the early signals in the wound response (Ca^2+^/ROS, apoptosis, MAPKs) was position-specific (see above). Instead, our data demonstrate that the number of apoptotic cells at the wounds site decreased upon Erk inhibition, but increased upon JNK inhibition (Fig. 4). The reduction of apoptotic cells upon Erk inhibition is consistent with observations in *Hydra* and in planarians (Chera et al., 2011; Owlarn et al., 2017) and further reports in which Erk activation in combination with high ROS levels results in the induction of apoptosis (Cagnol and Chambard, 2010). Although JNK activity has been frequently associated with the induction apoptosis, our data indicate non-apoptotic roles of JNK upon injury. This is consistent with the literature since JNK has been shown to have both pro-apoptotic and anti-apoptotic functions in various cell and tissue contexts (Krilleke et al., 2003; Liu and Lin, 2005).

### From injury to patterning

An important question in the field of regenerative biology is how does injury trigger pattern formation to replace lost structures. Our data have revealed an unexpected complexity between the early proteomic responses and the first transcriptional activation of the patterning GRNs. Previous work has revealed that nine out of eleven Wnt ligands and β-Catenin are activated in the early *Hydra* head organizer (Gufler et al., 2018; Hobmayer et al., 2000; Lengfeld et al., 2009; Petersen et al., 2015; Philipp et al., 2009; Technau et al., 2000). It was therefore striking that two of the “earliest” and one of the “latest” Wnt genes (Lengfeld et al., 2009) including β-catenin and the highly position specific target gene brachyury were not only activated during head regeneration, but also during foot regeneration (Fig. 6A) (Gufler et al., 2018). Since there is a <u>position-independent</u> activation of Wnt signaling, the previously proposed model that an immediate initial asymmetry in the expression of β-catenin and canonical Wnt signaling (Chera et al., 2009; Hobmayer et al., 2000) is difficult to explain, and it raises the question of what factors are initially responsible for relaying the positional information.

Our data reveal an unexpected complexity of MAPK and Wnt interactions. When we tested the influence of ERK, p38 or JNK activation on the expression of *Wnt3* and *Wnt910c*, we also found differences on Wnt activation in head and foot regenerates (Fig. 6 B-C). The inhibition of Erk caused a down-regulation of *Wnt3* in head-regenerates and a global reduction of *Wnt7* and *Wnt9/10c* expression, which is in accordance with the discovery that *Wnt1* expression is diminished upon abolished Erk activity (Owlarn et al., 2017). However, with respect to Wnt3 our data are only partly in line with the observation of an activating effect of ERK on Wnts irrespectively of position and wound type (Owlarn et al., 2017)(Reddien 2015). Conversely, the inhibition of p38 and JNK caused the upregulation of *Wnt3*, but a strong downregulation of *Wnt9/10c*. The antagonistic effects of p38/JNK on different Wnts was unexpected, but suggests a primary function of *Wnt9/10c* in the initiation of regeneration as polyps inhibited in stress-induced MAPK did not show any signs of regeneration (Fig. 1D) despite excessive *Wnt3* expression levels. We therefore propose *Wnt3* being a master regulator of patterning rather than a general initiator of regeneration although further experiments are required to validate that hypothesis. *Wnt7* had an intermediate pattern for ERK and p38/JNK in head and foot regenerates. Interestingly, qRT-PCR experiments also indicated a transition to position specific gene expression at 6 hpa, which is indicated by a drop of otherwise head specific Wnts in the foot regenerating animals (Fig. 6A). Remarkably, effects of MAPK inhibition became less or even non-significant at this time point. These data further support the hypothesis that MAPKs are part of a generic wound response initially regulating Wnt expression which later on becomes displaced by position specific cues that eventually regulate morphogenesis/patterning. This hypothesis is strongly supported by the work of Cazet & Juliano (back-to-back-submission) who found a general transcriptional response at 3 hpa regardless of the tissue context that becomes position specific at 8 hpa, thereby driving regeneration by positional cues. They further hypothesize that initial steps of Wnt expression are driven by bZIPs that are known targets of Erk and stress induced MAPK signaling pathways presented in this study.

### ATF/CREB mediated transcriptional activation of Wnts

Our *in-silico* analysis revealed a family of basic leucine zipper motif transcription factors, the activating transcription factors (ATFs), as potential candidates for the antagonistic effects of stress induced MAPKs on *Wnt* expression levels. While ATF2 has been reported to activate expression of target genes, ATF3 homodimers act as repressors (Chen et al., 1994; Vinson et al., 2002). Conversely, *Wnt9/10c* did not reveal ATF3 binding sites and accordingly showed a strong down-regulation upon inhibition of stress induced MAPK, which might be a consequence of absent ATF2 phosphorylation. Moreover, ATF3 is linked to cell survival, epithelial to mesenchymal transition, cell cycle progression, and stress responses (Yin et al., 2010). These pathways are also indispensable for proper regeneration (Tanaka et al., 2011) and it has been recently shown that ATF3 is also acting in the control of JNK signaling during *Drosophila* intestinal regeneration (Zhou et al., 2017). This points toward an important and promising role of ATF/CREB mediated transcriptional activation of *Wnts* in *Hydra*, particularly of ATF3.

In summary our data indicate an intimate link of the injury response and Wnt signaling. Our findings not only stress the importance of Wnt activation during regeneration, they also point to chronic injuries (e.g. viral hepatitis or excess alcohol consumption) where on top of an already existing genetic predisposition an injury-dependent increase in the risk of tumorigenesis was found (Arwert et al., 2012). Injured epithelia show increased activation of inflammatory signals, RTK and Wnt signaling to initiate healing by stem cell activation, proliferation, and ECM remodeling, so do cancer initiating cells to promote tumorigenesis (Arwert et al., 2012; Sundaram et al., 2017). Thus, our work on the elucidation how the injury response culminates into Wnt driven patterning may contribute to both the field of regenerative biology and cancer research (Gorlach et al., 2015; Romero et al., 2018).

## Acknowledgments

We thank Celina Juliano, Jack Cazet and Suat Özbek for critically reading the manuscript, Anna Marciniak, Moritz Merker, Mark Lommel, Sergio P. Acebrón, and Hendrik Petersen for helpful discussions, and Bruno Gideon Bergheim for support of bioinformatic work. The S65 and S66 strains with the Act hyGCaMP reporter were established by Jana Schlüter and Jan-Marek Weislogel and kindly provided by Hilmar Bading. The Funding: This work was supported by grants to T.W.H. (DFG-SFB 872-A1-2/3, DFG-SFB 1324-A5-1).

## Materials and Methods

### Hydra Culture

Polyps were kept in *Hydra* medium (1 mM NaHCO3; 0.1mM KCl; 0.1mM MgCl2; 1mM CaCl2; 1mM Tris pH 6.9,) at 18° C. Animals were fed three times a week with *Artemia salina* nauplii. The medium was exchanged daily. The strain *Hydra vulgaris* AEP was used for all experiments if not otherwise indicated. Animals were starved for 24 h prior to the performance of experiments.

### Transgenic HyGCaMP reporter animals

A PCR fragment containing 5’NheI -BglII -EcoRI 3’ restriction site was inserted into the hoT G actin EGFP vector, which yielded in the destruction of the 5’XbaI restriction site (Wittlieb et al., 2006). The vector containing GCaMP NLS (Ohkura et al., 2005) was optimized for *Hydra* codon usage with a BglII restriction site next to the NLS and synthesized by the Operon Company. HyGCaMP was amplified with primers containing NheI and EcoRI restriction sites (fw: 5’AA*AGATCT*AGCTTCGCTAGCATGG 3’; rv: 5’GT*GAATTC*TTAGGTGGATCTCTTAG 3’). After purification, HyGCaMP NLS was digested with BglII and EcoRI (NEB), which resulted in a fragment lacking the NLS. The same restriction enzymes were used for the hoT G actin ΔXbaI vector. After purification of both fragments, hoT G actin ΔXbaI and HyGCaMP ΔNLS were ligated using T4 ligase (Promega) according to the manufacturer’s instructions. The vector was used for microinjection of *Hydra vulgaris* AEP eggs at the 2-8 cell stage as described previously (Wittlieb et al., 2006). Fully transgenic animals were obtained by random budding.

Transgenic animals were either poked with a blunt glass pipette to obtain fully contracted polyps or cut into halves. The signal from the cytoplasmic HyGCaMP was imaged with a Nikon SMZ-25 stereoscope by taking movies with one picture per second. The background was subtracted from each picture and fluorescence was measured by a fixed region of interest at the area with initially highest signal intensity. Most distal from the emanating signal, a second ROI was used to measure the allover signal noise from the animal. Measured intensity upon stimulus was normalized to the tissue’s noise. Fold change was calculated using the signal of the unstimulated polyp as baseline value. Images were processed and measured using Fiji and results were visualized using GraphPad Prism 8.0.

### Inhibitor treatments

Animals were pre-treated with inhibitors for 1.5 h prior to the start of the respective experiment.

25µM U0126 (ERK inhibition), 50µM SB 203580 (p38 inhibition), 1µM SP600125 (JNK inhibition) were diluted in *Hydra* medium. All reagents were purchased from Cell Signaling Technology. For calcium experiments, 2µM A23187 (Sigma Aldrich) in *Hydra* medium were used if not otherwise indicated. 50µM BAPTA and 0. 5mM EGTA were used in calcium-free *Hydra* medium. Both reagents were purchased from Sigma Aldrich. For redox experiments, 100µM freshly purchased reduced glutathione (Merck Millipore) and 500µM H_2_O_2_ (Sigma Aldrich) were diluted in *Hydra* medium.

### Regeneration assay

Animals were bisected at 70% and 30% body length for head and foot regenerates respectively. Head regeneration was considered to be normal, when a hypostome and tentacle buds were detectable 3 dpa. Foot regeneration was considered to be normal, when differentiated tissue was detectable via peroxidase stain. The staining procedure was executed as previously described by Hoffmeister and colleagues (Hoffmeister and Schaller, 1985). For the staining reaction, TMB reagent (Sigma Aldrich) and H_2_O_2_ were diluted in 10mM citrate buffer pH 6.0. For ligature of *Hydra* tissue, the experiment was performed by carefully tying off the head using a human hair as described previously (Newman, 1974).

### Immunofluorescence analysis

Animals were bisected and relaxed in 2% urethane / *Hydra* medium for 1.5 minutes at 30 min post injury if not otherwise indicated. Polyps were shortly rinsed in 4% formaldehyde/ PBS-T (1x PBS/ 0.1% v/v Tween-20; 10xPBS: 8%w/v NaCl, 0.2 %w/v KCl, 2.68%w/vNa_2_HPO_4_, 0.24 %w/v KH_2_PO_4_, pH 7.4). Fixation solution was replaced by fresh solution and animals were fixed at 4°C overnight. On the next day, polyps were washed 3 times in TBS-T and subsequently solubilized in TBS/0.1% v/v Triton-X 100. Solubilization buffer was removed by three washes in TBS-T (1x TBS/ 0.1% v/v Tween-20; 10xTBS: 0.2M Tris pH 7.4, 1.5M NaCl). To recover antigens, Tris based retrieval was performed as described by Inoue and colleagues (Inoue and Wittbrodt, 2011). Animals were incubated in TBS-T for 10 min and subsequently transferred to antibody solution using a 1:100 dilution in blocking buffer (1% v/w BSA, TBS-T, 5mM EDTA pH 8.0, 1mM EGTA pH 8.0) at 4°C overnight. Excess of antibody was removed by three washes in TBS-T followed by incubation with the secondary antibody (Goat anti-Rabbit IgG Alexa Flour 488; # A-11034; Thermo Fisher Scientific) at a 1:500 dilution in blocking buffer in dark for 2.5h at RT. Excess of antibody was removed by three washes in TBS-T, where DAPI (Roche) was added 1:1000 to the second washing step. Polyps were mounted in Mowiol 4-88 (Roth). First antibodies used in this study were all purchased from Cell Signaling Technology and can be found under the following product numbers: Phospho-ERK1/2: #4370; Phospho-p38: #4511; Phospho-JNK:#4668. Animals were imaged with Nikon Eclipse 80i or Nikon A1 confocal microscope. Pictures were processed with Fiji and Adobe Photoshop.

### CellRox Assay

Animals were incubated with 5µM of Molecular Probes™ CellRox Green Reagent (Thermo Fisher Scientific) in *Hydra* medium for 12 h at 18°C. Animals were bisected or tied and subsequently transferred into fresh *Hydra* medium. Reaction was stopped after 20 minutes by incubating polyps in 2% urethane/ *Hydra* medium followed by fixation in 4% formaldehyde/ PBST for 20 min at 4°C. Polyps were then mounted in Mowiol 4-88 (Roth). Animals were imaged with Nikon Eclipse 80i. Pictures were processed with Fiji and Adobe Photoshop.

### Western blot

Animals were cut 3 times along the body column. For comparability, animals that have been treated in parallel to tied animals, polyps were cut only once. Whole tissue lysates have been taken up in lysis buffer (4% w/v SDS; 0.5% w/v sodium deoxycholate; 20 mM Tris pH 7.5) and heated at 95°C at 800rpm in a heating block for 10min. After centrifugation at 15 000 xg for 10min, protein concentration was determined using the enhanced protocol of the Pierce BCA assay kit (Thermo Fisher) according to the manufacturer’s instructions. 10µg protein was mixed with 5x sample buffer (500 mM Tris pH6.8, 50% v/v glycerol; 10% w/v SDS, 0.1% w/v bromophenol blue, 10% beta-mercaptoethanol), boiled at 95°C for 10 min and subsequently applied for discontinuous SDS gel electrophoresis using a 12% separating gel (10x running buffer: 1.92 M glycine; 10% w/v SDS; 0.25 M Tris). Gels were run at 20 mA per gel. Proteins were then transferred to nitrocellulose membranes (Amersham; 0.45 µM NC; GE Healthcare) at 360 mA for one 1 h or 30 mA overnight in blotting buffer (10% v/v 10x running buffer; 20% v/v methanol). Membranes were blocked in blocking buffer (5% v/w BSA, TBS-T, 5mM EDTA pH 8.0, 1mM EGTA pH 8.0). First antibodies were diluted 1:1000 in blocking buffer containing 1% w/v BSA and incubated at 4°C overnight. On the next day, membranes were washed three times in TBS-T for 10 min each and incubated with the secondary antibody diluted 1:5000 (loading control) or 1:2000 (MAPK) in blocking solution for 2h at RT. Membranes were washed three times in TBS-T for 10 min each. Light reaction was initiated by applying ECL1 and ECL2 (Thermo Fisher) to the membranes and imaged with Chemostar ECL imager (Intas). For detection of phospho-specific MAPKs, the same antibodies were used as detailed in the immunofluorescence section. As loading control, mouse anti-alpha-tubulin (#T6199; Sigma Aldrich) was used. For the detection of total protein amounts the following antibodies were purchased from Cell Signaling Technology: ERK1/2 : #4695; p38: #9212; JNK: #9252. As secondary antibodies, goat anti-mouse HRP conjugate (Jackson Immuno Research) and goat anti-rabbit HRP conjugate (# 7074; Cell Signaling Technology) were used. For quantification, obtained bands were measured with the gel analyzer tool of Fiji. The values of the measured areas were normalized to the obtained ones of the loading control. Fold change was calculated using the t_0_ value as reference.

### TUNEL Assay

In-*situ cell* Death Detection kit Fluorescein (Roche/ Sigma-Aldrich) was used to detect apoptotic cells. Animals were relaxed in 2% urethane / *Hydra* medium for 1.5 minutes at 2 h post injury if not otherwise indicated. Animals were shortly rinsed in 4% formaldehyde/PBS-T and transferred to fresh formaldehyde solution. After overnight fixation at 4°C, animals were washed 3 times in TBS-T for 10 minutes each followed by 3 washes in TBS/ 0.1%Triton for 10 minutes and thereafter, 3 times in TBS-T for 10 minutes each. Animals were equilibrated in 10mM citrate, pH 6.0 / 0.1% Tween-20 for 15 minutes at room temperature. After, 225μl label solution and 25μl enzyme solution were added to the animals and incubated for 90 minutes at 37°C in dark. Afterwards, animals were washed in 1x TBS-T for 10 minutes. In the second washing step, DAPI (Roche/Sigma Aldrich) was added to wash and incubated for 10 minutes. After, animals were washed a third time in TBS-T for 10 minutes and subsequently mounted in Mowiol 4-88 (Roth). Animals were imaged with Nikon Eclipse 80i. Pictures were processed with Fiji.

### RNA isolation and cDNA synthesis

Polyps were pre-incubated in *Hydra* medium supplemented with DMSO or inhibitor for one hour and subsequently bisected at 50% body length. Regenerating tips were dissolved in 1mL Trizol (Thermo Fisher) and 0.2mL chloroform (Sigma Aldrich). After centrifugation at 12,000 x g at 4°C, upper phase was transferred into a fresh tube and mixed with chloroform:isoamylalcohol at a ratio of 24:1. Samples were spun down again as described above and upper phase was transferred into a fresh tube again, where RNA was precipitated with 0.8V pure isopropanol. After centrifugation, the pellet was washed once in 75% v/v ethanol and afterwards dried by air before it was taken up in nuclease-free water. RNA was digested with 1.5 U DNaseI (Roche/Sigma Aldrich) and inactivated according to the manufacturer’s instructions. The quality of RNA was examined by agarose gel electrophoresis and NanoDrop. cDNA was transcribed using 1µg RNA and sensiFAST cDNA Synthesis Kit (Bioline) following the instructions of the manufacturer.

### Quantitative RT-PCR

cDNA was diluted 1:10 in nuclease-free water and subsequently applied for qRT-PCR using the SensiFast™SYBR HiROX Kit (Bioline) according to the manufacturer’s instructions. Data were normalized to GAPDH and evaluated using the ΔΔCt method. Results were visualized by GraphPad Prism 8 and tested for significance using t-test and Kruskal Wallis test. *Wnt3 fw*: 5’ATTACAACAGCCAGCAGAGAAAG 3’; *rv*: 5’TTATCGCAACGACAGTGGAC 3’; *Wnt7 fw*: 5’CGGCAATAATTGAATGAACAAAT 3’, *rv*: 5’ACCTGGTTGGCAATCACAAT 3’; *Wnt9/10c fw*: TGCTCTATAGTGTCTGGTTCTTTACT 3’, *rv*: 5’ TGAGCTAAACTTGCCGAGAATA 3’; *GAPDH fw*: 5’GAGCATCCTGATATTGAAATTGTTC 3’, *rv*: CATGGTATTTCTTTTGGGTTTCTAA 3’

### Wound induced budding

Budless animals were labelled with an injection needle filled with fluorescent microspheres (Flouresbrite YG Microspheres 1.00µm; Polysciences) that have been diluted 1:10 in *Hydra* medium prior to use. 2h post labelling, one half of the population was cut in the budding region at the side of the label, while the other half remained unaffected. Buds relative to the label were counted 3 days post labeling. Statistical relevance was evaluated using logistic regression analysis.

### Promotor prediction

For the prediction of promotors, the published sequence for *Wnt3* has been used (Nakamura et al., 2011), while the other sequences were retrieved by blasting the open reading frame of the *Wnt7* and *Wnt9/10c* against the *Hydra* 2.0 Genome. The first 3-2 kb downstream of the transcription start site of *Wnt 7* (ID: Sc4wPfr_1214), *Wnt9/10c* (ID: Sc4wPfr_224.1), and Wnt3 were used for prediction of potential transcription factor binding sites using PROMO3 with a 5% maximum matrix dissimilarity rate (Farre et al., 2003; Messeguer et al., 2002). Motifs were taken into consideration when RE equally < 0.1. This cut-off was deduced from values obtained for TCF sites of the *Wnt3* promotor since the RE value correlated with experimentally determined functionality.

## Notes

### Competing Interest Statement

The authors have declared no competing interest.

