## Supplemental Figures for "MAPK signaling links the injury response to Wnt-regulated patterning in *Hydra* regeneration"

### Supplementary Figures

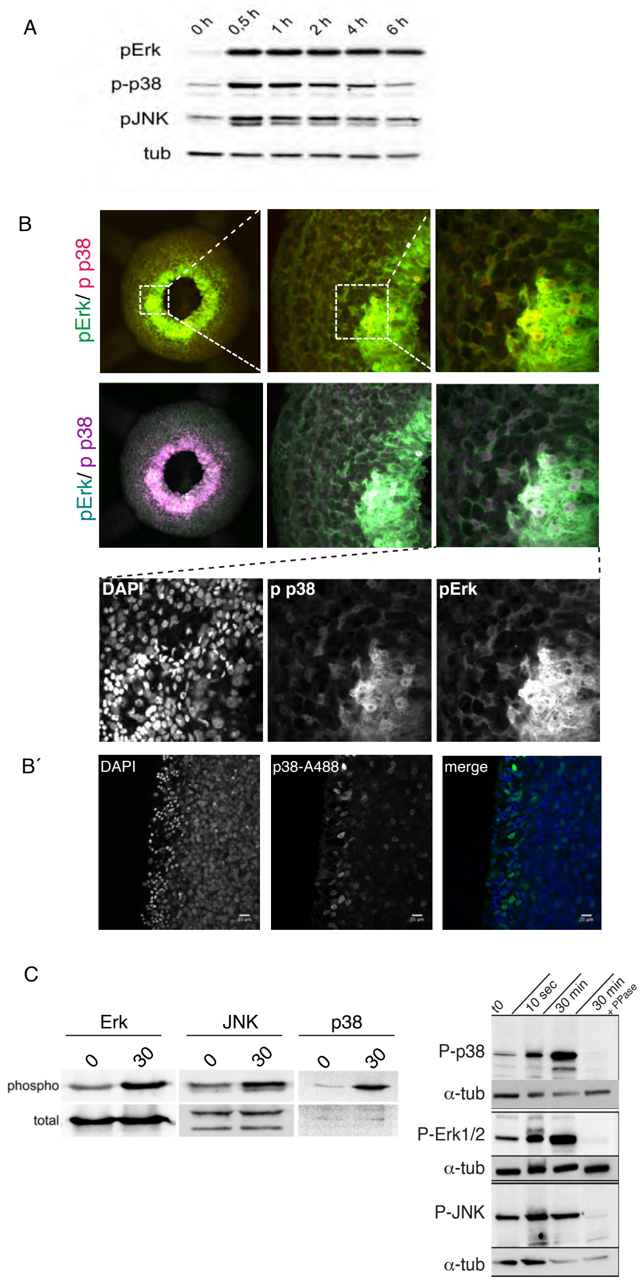


**Figure S1. MAPKs are spatially and temporally distinct. A)** While ERK shows elevated phosphorylation levels after 6hpi, levels of pJNK and p-p38 drop much faster. **B)** Confocal images of standard phospho-ERK antibody and p-p38 antibody already conjugated to an Alexa-flour-597 antibody (Cell Signaling Technology **#8632).** Phosphorylated p38 resides in nuclei of pERK positive nuclei. **B')** Confocal images of *Hydra* demonstrating that p-p38 localizes to weakly stained nuclei. **C)** Western blot analysis of ERK, p38, and JNK demonstrating that after 30 minutes post injury, only phosphorylation levels increase while total protein levels remain unaffected. **C')** Treatment with alkaline and lambda phosphatase prior to incubation with the first antibody resulted in absent bands, demonstrating the phospho-specificity of the phospho-antibodies.

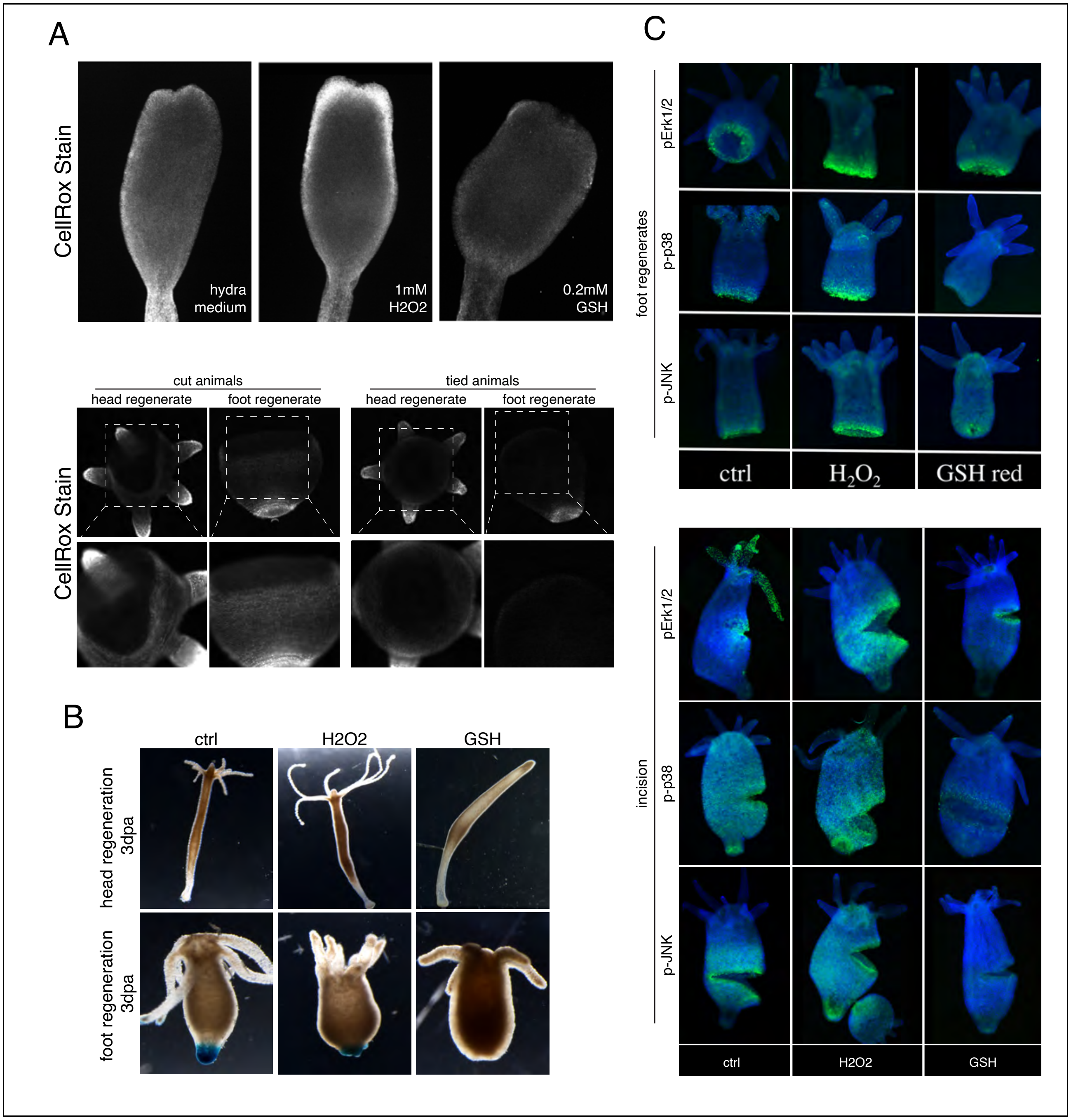


**Figure S2. Activation of MAPKs is ROS-dependent. A)** CellRox stain results in an increase of signal intensity when cut polyps were incubated with $H_2O_2$, while reduced glutathione did not result in any detectable signal. CellRox positive cells were detectable in both, head and foot regenerates, while ligature of the tissue did not reveal the generation of ROS. **B)** Representative pictures of head and foot regenerating polyps upon $H_2O_2$and GSH exposure. Differentiated foot tissue was visualized by peroxidase color reaction. **C)** Increased levels of MAPK upon $H_2O_2$ exposure are likewise detectable in foot regenerates and upon incision, while GSH yields in a reduction.

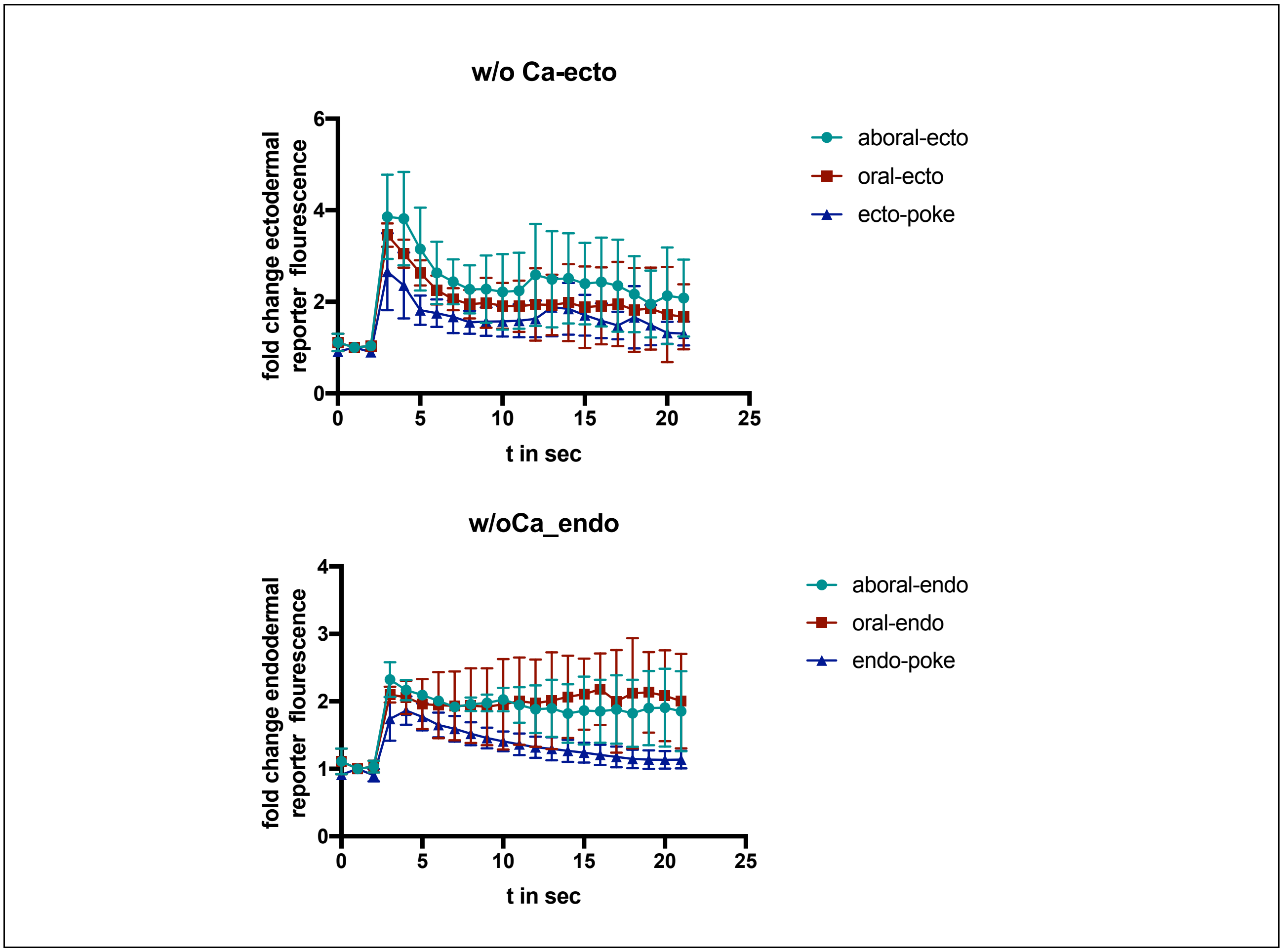


**Figure S3. Reporter activity of GCaMP-transgenic animals in calcium-free medium.** Background was subtracted from images and signal generated by poking or injury was normalized to the intrinsic signal most distal from the origin of the reporter signal. Compared to poked animals, injured animals exhibited a stronger sustained signal.

**Additional Supplementary Movies**

**S3_movie1**. Ectodermal calcium reporter animals upon poking
**S3_movie2**.Ectodermal calcium reporter animals upon amputation
**S3_movie3**. Ectodermal calcium reporter animals upon poking in calcium-free medium
**S3_movie4**.Ectodermal calcium reporter animals upon amputation in calcium-free medium
**S3_movie5**. Endodermal calcium reporter animals upon poking
**S3_movie6**.Endodermal calcium reporter animals upon amputation
**S3_movie7**. Endodermal calcium reporter animals upon poking in calcium-free medium
**S3_movie8**.Endodermal calcium reporter animals upon amputation in calcium-free medium

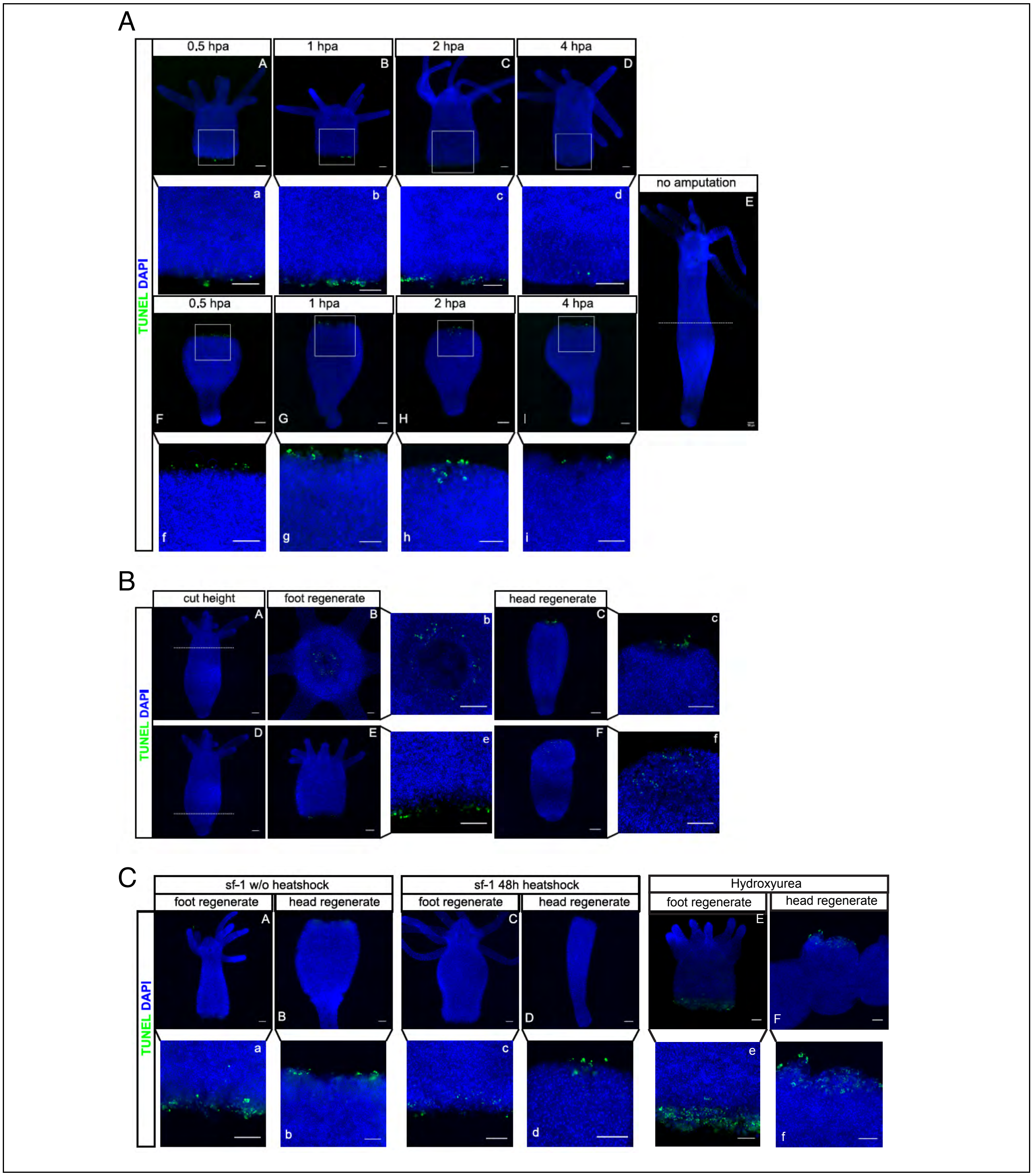


**Figure S4. Apoptosis upon injury. A)** TUNEL positive cells were detectable in head and foot regenerates with a maximum between 1-2 hpa. **B)** Frequency of apoptotic cells was independent from cut height. **C)** Loss of i-cells was either induced by heat shock of sf-1 animals or by incubating animals with hydroxyurea followed by a three days chase. For each condition, comparable numbers of TUNEL-positive cells were detectable with the exception of hydroxyurea-treated animals that showed elevated frequencies. Scale represents 100µm.

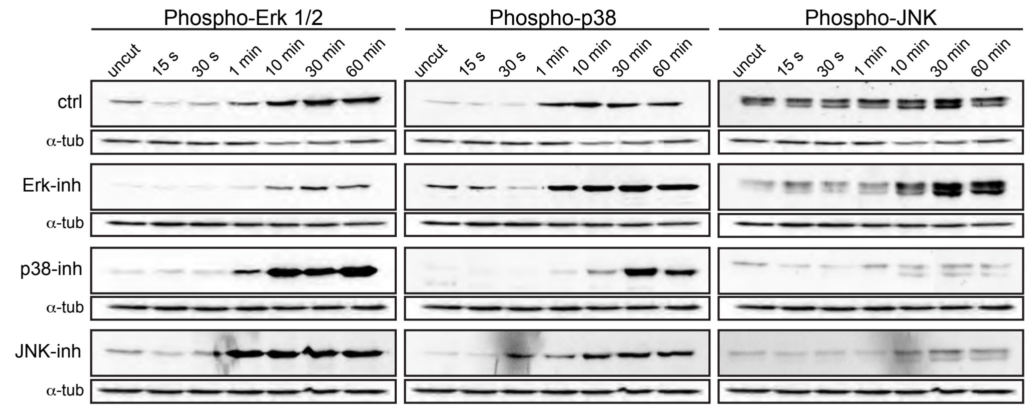


**Figure S5. MAPKs underlie extensive crosstalk.** Western blot analysis of pERK, p-p38, and pJNK upon pharmacological inhibition of the mentioned MAPKs.

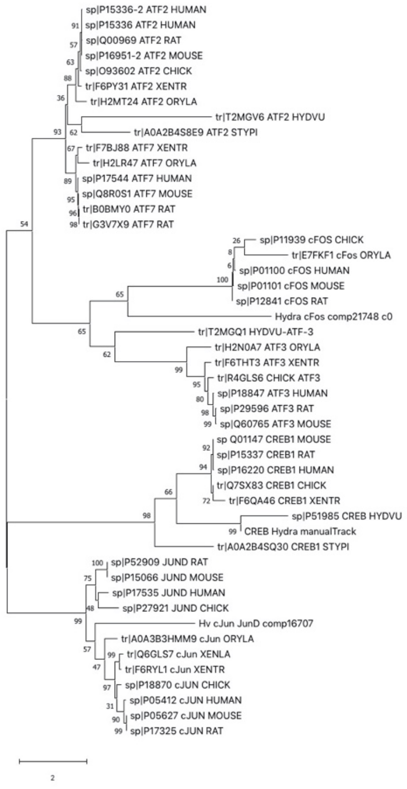


**Figure S6. ATF phylogeny. A)** Phylogenetic analysis of ATF2 and ATF3 to validate the presence of these homologues in *Hydra*. **B)** Alignment of protein sequences used for phylogenetic analysis (see next page).

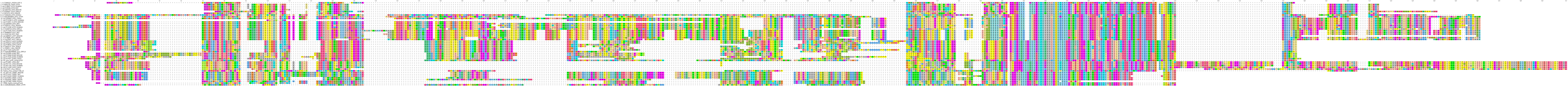
